# Inhibition of hyperactive cyclin dependent kinase 5/p25 is protective in the 6-hydroxydopamine model of Parkinson’s disease

**DOI:** 10.1101/2022.10.03.510400

**Authors:** Ashley Bernardo, Niranjana Amin, BK Binukumar, Harish Pant, Ram Mishra

## Abstract

**Background:** Cyclin-dependent kinase 5 (CDK5) is a multifunctional enzyme involved in neuronal development, maturation and survival. CDK5 activity is tightly regulated by association with regulatory proteins p35 and p39. Upon neuronal insults, increased intracellular calcium activates calpain, cleaving p35 into p25, which has a higher affinity for CDK5. p25 hyperactivates CDK5, initiating apoptotic cascades that lead to significant dopaminergic (DAergic) loss that can leads to neurodegenerative disorders, such as Parkinson’s disease (PD).

**Objective:** This study investigates hyperactivation of CDK5/p25 in the 6-hydroxydopamine (6-OHDA) rat model of PD and specific inhibition of CDK5/p25 by truncated peptide 5 (TP5). TP5 was investigated for amelioration of 6-OHDA induced behaviour impairments and significant protection of dopamine neurons through tyrosine hydroxylase (TH).

**Methods:** 6-OHDA induced motor impairments and reduced TH. Motor assessments included locomotor activity, beam transversal, fixed speed rotarod and amphetamine-induced rotations. Immunohistochemistry investigated DAergic neurodegeneration using TH levels and immunoprecipitation and assay investigated CDK5 activity.

**Results:** Pre-administration of TP5 maintained locomotor activity, preserved beam transversal scores, protected motor coordination and attenuated amphetamine induced rotations in 6-OHDA lesioned rats, all indicative of neuroprotection by TP5. 6-OHDA without pretreatment of TP increased CDK5 activation. CDK5 activity in TP5+6-OHDA animals was not significantly different from artificial cerebrospinal fluid (aCSF) treated sham surgery controls. Immunohistochemistry revealed significant TH protection within the substantia nigra (SN) of TP5 pretreated animals.

**Conclusions:** 6-OHDA increases CDK5 activity. Hyperactive CDK5/p25 inhibition in the 6-OHDA model has neuroprotective capability, protecting against the development of a toxin-based induction of PD-like motor phenotypes and pathology. This supports CDK5/p25 specific inhibition as a target for further neuroprotective therapeutic development.

## 1.0 Introduction

Parkinson’s disease (PD) is a debilitating neurodegenerative disorder characterized by the degeneration of dopamine neurons within the substantia nigra and nigrostriatal circuitry (Radhakrishnan and Goyal, 2018). Resulting motor symptoms include tremors, bradykinesia and muscle rigidity while non-motor symptoms include memory loss, speech impairments, depression and anxiety (Galvan and Wichmann, 2008; Magrinelli et al., 2016; Schapira et al., 2017). The “gold standard” treatment is dopamine replacement therapy levodopa (L-dopa). With disease progression, L-dopa loses efficiency and coincides with other motor and non-motor side effects (Rascol et al., 2003). L-dopa is also metabolized by gut microbiota reducing its bioavailability (van Kessel et al., 2019). Progressive dopaminergic degeneration diminishes treatment response and therapeutics struggle to provide neuroprotection. It is hypothesized that dopaminergic depletion occurs from increased oxidative stress, endoplasmic reticulum stress and subsequent hyperactivation cyclin-dependent kinase 5 (CDK5) activity (Cruz et al., 2003; Qu et al., 2007; Schapira, 2008; Blesa et al., 2015).

CDK5 is a neuron-associated serine/threonine kinase involved in neuronal migration, differentiation, synapse development, synapse function and apoptosis (McLinden et al., 2012). Binding of CDK5 to specific promoters (p35, p25 (a c-terminus fragment of p35) or p39) determine anti-apoptotic or pro-apoptotic downstream effects of CDK5 (Hisanaga and Endo, 2010; McLinden et al., 2012). The role of p35 in neuronal survival is well established by inhibiting c-Jun N-terminal kinase (JNK) 3 and activating the neuregulin/ phosphatidylinositol 3-kinase (PI3K)/Akt pathway (Li et al., 2002, 2003). During physiological stress, p35 is cleaved by the calcium dependent enzyme calpain into p25. Cleavage converts pro-survival p35 into shorter, pro-apoptotic p25 fragment (Kusakawa et al., 2000). CDK5 bound to p25 phosphorylates nuclear proteins inducing pro-apoptotic events, damages DNA and arrests the cell cycle. In disease states like PD, increased presence, and longer half-life of p25 hyperactivates CDK5. The CDK5/p25 complex increases neuronal degradation by inhibiting paired-related homeodomain protein 2 (Prx2) (a reactive oxygen species (ROS) scavenger). This results in accumulation of ROS and oxidative stress (Qu et al., 2007). CDK5 has been implicated through increased calpain and CDK5 in several neurodegenerative diseases such as Alzheimer’s disease (AD) and PD through post-mortem studies and pre-clinical models both reviewed well in Wilkaniec et al (Wilkaniec et al., 2018).Treatments attempting to inhibit elevated CDK5 activity have so far proved ineffective because they inhibit normal pro-survival p35-bound CDK5 (Binukumar et al., 2015; Siklos et al., 2015).

Novel peptide, truncated peptide 5 (TP5) is a 24-amino acid peptide designed based on the structure of p35 (Binukumar et al., 2015) under patent #US8597660B2. TP5 inhibits CDK5/p25 specifically while having minimal effects on the endogenous CDK5/p35 complex (Zheng et al., 2005, 2010). TP5 has been investigated for neuroprotective properties in models of neurodegenerative disorders such as AD (decreases neurofibrillary tangles, inflammation and amyloid plaques) and ischemic stroke (Shukla et al., 2013; Ji et al., 2017). Neuroprotective properties have been documented using TP5 in 1–methyl-4-phenyl-1,2,3,6-tetrahydropyridine (MPTP/MPP(+)) animal and cell culture models of PD (Binukumar et al., 2014, 2015). Functional neuroprotection related to PD is important to evaluate during investigation of novel potential therapeutics. We hypothesized that 6-OHDA will increase CDK5/p25 activity and TP5 will have neuroprotective effects by reducing CDK5/p25 hyperactivation. This study considered a battery of motor function and coordination tasks to investigate if neuroprotection exhibited by TP5 in the substantia nigra (SN) extends to PD-like motor impairments developed in the 6-hydroxydopamine (6-OHDA) model of PD. We further demonstrate neuroprotective properties of TP5 through investigation of CDK5 levels and tyrosine hydroxylase (TH).

## 2.0 Experimental Procedures

### 2.1 Animals

Male, Sprague Dawley rats (N=29), 8 weeks old, weighing 250-300g were obtained from Charles River Laboratories (Quebec, Canada). Upon arrival all animals were handled for 7 days prior to initiation of experiments. This habituated animals to experimenters and reduces variability induced by handling anxiety. Animals were individually housed at McMaster University Central Animal Facility at 21°C, on a 12-hour dark/light cycle and on a food restricted diet to maintain 95% of their free feeding weight. Food restriction was required as beam walk used food as an incentive to motivate animals into performing the test and overcomes inconsistent intervals between an animals last feeding time and testing time if given food access *ad libitum*. Rats received water *ad libitum*, and all experimental procedures were completed in accordance with the Animal Research Ethics Board and Canadian Council on Animal Care. 14 animals were used for behavioral work. 2 of the original 16 were euthanized based on endpoint criteria of significant weight loss and dehydration, one from ScP+6-OHDA and one from aCSF+6-OHDA. Of these, 3 from each group were used for immunohistochemistry, 3 were used for CDK5 activity in the aCSF+6-OHDA group and one in each of the other groups. An additional 13 animals were purchased for CDK5 activation studies (4 control, 3 aCSF+6-OHDA, 3 ScP+6-OHDA and 3 TP5+6-OHDA). The MPTP and 6-OHDA neurotoxin models of PD have overlapping mechanisms to induce cell death, motor deficits and they have similar induction rates. Therefore, we can use the previous study by Binukumar et al. to evaluate sample size. Previous work with 8 animals showed very low variability in both CDK5 activity and behavioural phenotypes such as locomotion, suggesting less animals were justified to be used in the present study. The present study n was also based on housing occupancy and behavioral testing feasibility. However, no specific power analysis was used to determine sample size.

### 2.2 Surgery

Animals were anesthetized using Isoflurane and oxygen. Animals were mounted on a stereotax and an incision was made along the skull. Bregma was located and stereotaxic coordinates were documented. At stereotaxic coordinates A/P: −5.3mm and M/L: −2.3mm in reference to bregma, from the Paxinos and Charles Watson’s rat brain atlas (2nd edition), a hole was drilled above the SN. A second hole was drilled close to the first hole and a jeweller’s screw was secured to act as an anchor for the permanent cannula. The permanent cannula (HRS Scientific, Quebec) was lowered into the substantia nigra (D/V: −7.3mm from brain surface) using the stereotax. Dental cement was used to attach the cannula to the jeweller’s screw. Once secured, a dust cap was inserted into the cannula to avoid clogging and prevent infection. The incision was closed, and animals were given 5 days to recover before infusions began.

### 2.3 Treatments

TP5 and Scrambled peptide (ScP) were synthesized by Peptide 2.0, a company that guarantees peptide purity (Chantilly VA, USA) (Binukumar et al., 2014, 2015; Cardone et al., 2016). 1mg of TP5 was dissolved in 150μL of sterile water and a cumulative total of 40μg in 6μL was administered to the substantianigra. The following sequences were used with bold indicating a TAT tag that allows the peptides to cross the blood brain barrier:

TP5: KEAFWDRCLSVINLMSSKMLQINA**YARAARRAARR**

ScP: GGGFWDRCLSGKGKMSSKGGGINA**YARAARRAARR**

While full pharmacokinetic studies have yet to be completed, preliminary work from our group has found the peptides to be highly stable on the basis of a Fluorescein isothiocyanate (FITC) tag attached to TP5 and ScP detected up to 1 week later within an *in vitro* setting. This data is unpublished. 6-OHDA was obtained from Sigma-Aldrich (Oakville, Ontario) and dissolved in 0.9% saline containing 0.2% ascorbic acid. 6-OHDA was kept on ice and away from light.

### 2.4 Infusions

Infusions commenced 5 days after surgical placement of cannulas. The infusion cannula was connected to a Hamilton syringe placed in a Harvard Apparatus PHD 2000 syringe pump via a short tube. The infusion cannula was checked for blockages, inserted into the permanent cannula (D/V: 7.8mm) and attached using the connector assembly (HRS scientific, Quebec). All animals received 3 infusions under anesthesia at an infusion rate of 1uL/min. Infusion one occurred 10 hours prior to 6-OHDA ensuring time for solution uptake by cells. Depending on treatment group animals received 4μL of aCSF, ScP or TP5 as infusion one. Infusion two (2μL of the same treatment previously received) occurred 1.5 hours prior to 6-OHDA insult. Infusion two increased the amount of treatment at the site of 6-OHDA administration to account for any degradation by peptidases. Third infusion was 8μg 6-OHDA in 3μL administered to all treatment groups other than control used in biochemical assessment, which received aCSF.

### 2.5 Behavioural testing

Animals were habituated, trained and baseline tested for motor tasks prior to surgery. No differences were found between groups prior to surgery. Behavioural testing was conducted 7 days after infusions. Experimental timeline is depicted in Figure 1. One behavioural test was completed per day for all animals in the experiment in the following order: beam walk, fixed-speed rotarod, locomotor activity and amphetamine induced rotations. For each test, 3 trials were performed per animal and averaged for presented results, with the exceptions of amphetamine induced rotations and locomotor activity. This is because multiple administrations of amphetamine produces mechanistic changes that are considered to induce a model of Schizophrenia. Locomotor activity was only assessed once after interventions as the novelty of the chamber is no longer present therefore animal exploration is less apparent.

**Figure 1.**
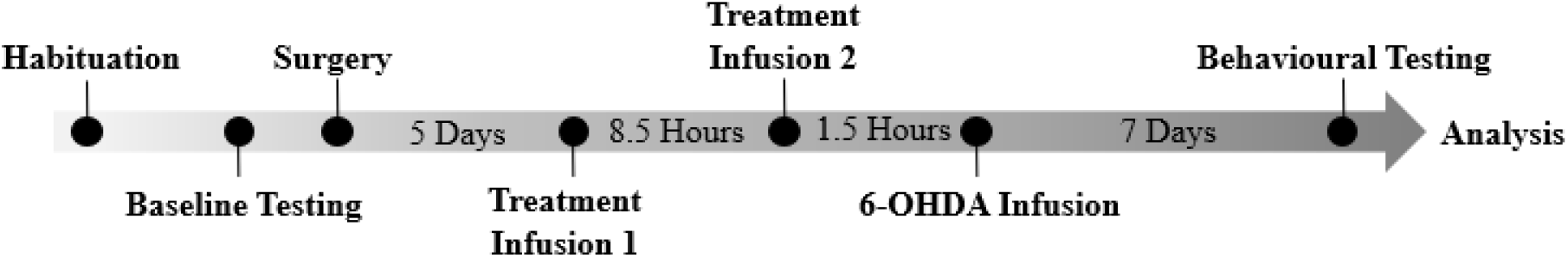
Experimental Timeline. Visual representation of the experimental treatment timeline and behavioural work. Important temporal elements depicting the amount of time between two events that are shown as black circles. Treatments given at infusion 1 and 2 were dependent on treatment group: aCSF+6-OHDA group received aCSF for infusions 1 and 2, ScP+6-OHDA animals received ScP for infusions 1 and 2 and the TP5+6-OHDA group received TP5 at infusions 1 and 2. All groups other than controls received 6-OHDA in infusion 3. Controls received aCSF at all three infusions.

### 2.6 Locomotor activity

Animals were placed in AccuScan computerized chambers (Accuscan Instruments, Columbus, OH). Chambers measured animals’ locomotor activity in the x and y planes using infrared beams evenly distributed front to back and left to right. When the animal travels, specific beams are broken, and VersaMax Analyser software identified the total distance travelled by the animal. Total distance travelled was recorded for all animals over a 2-hour period.

### 2.7 Beam walk

Beam walk measured motor coordination and balance (Allbutt and Henderson, 2007). A flat surface starting platform and 20cmx20cmx20cm enclosed finishing platform were 60cm above the table and 1 meter apart. Animals were trained until they could easily transverse a 1.5cm wide beam using a Kellogg’s Froot Loop as incentive at the finishing platform. Parameters recorded for beam walk by 2 blinded experimenters were 1. time to initiate transversal, 2. time to transverse the beam and 3. number of hindlimb slips. Training was considered complete when animals obtained a score of <1 meaning on average they could initiate transversal in < 30 seconds and cross in <3 seconds with 0 slips. The following scoring system, based on Urakawa et al. 2007 (Urakawa et al., 2007), was used to analyse the testing phase of beam walk: 0= transverses with no difficulty (time to cross <3 seconds, 0 slips), 1=transverses with little difficulty (>3 seconds but <5 seconds to transverse, 0 slips), 2=walking disability but can transverse (<3 seconds to transverse with 1 or more hindlimb slips), 3=considerable time to transverse the beam (>3 seconds to transverse with 1 or more slips), 4=unable to transverse beam (>3 seconds to transverse but unable to reach the end (falls), 5=unable to transverse and remains on starting platform (>30 seconds to initiate and unable to reach end platform (falls)). 3 trials per animal were completed and their scores were averaged.

### 2.8 Fixed speed rotarod

The fixed speed rotarod (FSRR) evaluates balance and sensorimotor coordination (Monville et al., 2006). The FSRR apparatus has a revolving 7cm in diameter, motor driven rod with an automated switch below. Animals were placed on the FSRR by the experimenter. Animals underwent three trials at 10rpm. The latency to fall off the rod in each trial was recorded by the switch below and averaged for analysis.

### 2.9 Amphetamine induced rotations

Amphetamine induced rotation is a phenomenon present in unilateral 6-OHDA lesion models of PD. Receptor supersensitivity in one hemisphere of the brain produces rotation after 6-OHDA induced damage and phenotypically evaluates dopamine loss (Ungerstedt and Arbuthnott, 1970; Mishra et al., 1974, 1980; Björklund and Dunnett, 2019). 2.5mg/kg of amphetamine was administered intraperitoneally, and animals were placed in a clear cylinder 31cm tall and 30cm in diameter. Animals were recorded over a 30-minute period. Ipsilateral and contralateral rotations (in reference to the permanent cannula) was analyzed from the last ten minutes of the video by a blind researcher. Ipsilateral minus contralateral rotations are presented here.

### 2.10 Tissue collection for CDK5 assay

Animals were anesthetized using isoflurane and subsequently decapitated. The whole brain was removed, the SN was dissected while on ice and flash frozen before storage at −80°C.

### 2.11 Tissue collection for immunohistochemistry

Animals were anesthetized using a ketamine (75mg/kg) and xylazine (10mg/kg) mixture and perfused with PBS then 4% paraformaldehyde (PFA). The brain was extracted and stored in 4% PFA overnight. The following day the brain was immersed in 10% sucrose for 24 hours followed by 30% sucrose. Each brain was flash frozen using 2-methyl butane and stored at −80°C. Chromium-gelatin-coated slides were prepared and 16μm thick serial, coronal sections were obtained of using a cryostat. Slides were stored at −80°C.

### 2.12 CDK5 immunoprecipitation and assay

Kinase assays were performed as described previously, with modification (Binukumar et al., 2014). Briefly, CDK5 was immunoprecipitated with polyclonal C8 antibody for 2 h at 4°C and protein A–Sepharose beads were used to isolate immunoglobulin. Immunoprecipitates were washed three times with lysis buffer and once with 1X kinase buffer. 1X kinase buffer contained 5 mM MOPS, pH 7.4, 2.5 mM β-glycerophosphate, 1 mM EGTA, 0.4 mM EDTA, and 5 mM MgCl_2_. Samples were added to the reaction mix containing kinase buffer, 50 μM ATP, 20 μg of histone H1, and 0.1mCi of [γ^32^P]ATP containing 0.1mm DTT and 1X Halt protease and phosphatase inhibitor (Thermo Fisher) and incubated at 30°C for 1 h. Reactions were stopped by adding Laemmli sample loading buffer, and samples were electrophoresed on 12% SDS–PAGE gels. Histone bands were visualized by Coomassie blue staining and gels were autoradiographed and were scanned on a PhosphorImager. Radioactive band density was analyzed using ImageJ software, and statistical analysis was performed.

### 2.13 Immunohistochemistry

Slides were washed 3 times in PBS, post fixed with 4% PFA for 10 minutes and washed 3 time with PBS. Slides were blocked using BSA, normal goat serum, 0.1% Triton X and PBS before incubation with anti-TH antibody (dilution 1:500) (Cedarlane, Ontario Cat.# CL8876AP, Lot#167621) overnight at 4°C. Specimens were washed in PBS and incubated for one hour at room temperature using Texas red goat anti-rabbit antibody (1:1000) (Invitrogen, Ontario). Coverslips were adhered using PureLong Gold mounting medium containing DAPI (Life Technologies). Sections were imaged using Leica DM6 B microscope and Leica Application Suite X (LAS X) software. Immunofluorescence was analyzed by ImageJ software and reported as integrated density. Integrated density was obtained after conversion to 8-bit images and background was subtracted. The wand tool was used to generate the region of interest by outlining the SN based on the fluorescence edges.

### 2.14 Statistical analysis

Behavioural and biochemical data were analysed using One Way-ANOVA with Tukey’s post hoc using GraphPad Prism software. Group was always considered the independent variable and significance was considered as *p<0.5, **p<0.01, ***p<0.001. All error bars are represented as standard error of the mean (SEM). Analysis was completed on all animals. If an outlier was detected by Grubbs Outlier test, we noted if its exclusion changes statistical significance.

## 3.0 Results

### 3.1 Locomotor activity in affected in 6-OHDA lesioned rats treated with aCSF, ScP or TP5

Locomotor activity was assessed through total distance travelled to distinguish the rodent equivalent phenotypic manifestation of rigidity, akinesia and bradykinesia. Accuscan technology found an effect of group (F13)=6.073, p=0.0167. Overall, TP5+6-OHDA animals travelled more than those receiving aCSF+6-OHDA. Higher total distance travelled found in TP5+6-OHDA treated animals compared to other groups, suggests pre-treatment with TP5 was able to protect the development of reduced motor activity inflicted by 6-OHDA. One outlier is detected in the aCSF group. With its exclusion there is no change to significance.

### 3.2 TP5 improves beam walk in 6-OHDA lesioned rats

Beam walk measured balance and coordination. Using a predetermined scale described in experimental procedures, beam walk considers animals’ ability to initiate movement onto the beam, speed of transversal and balance abnormalities through hindlimb slips off the beam. Animals pre-treated with TP5 before 6-OHDA achieved a score indicative of little difficulty crossing the beam. A significant effect of group was found using One Way ANOVA (F(10)=9.826, p=0.0036). Tukey’s post hoc testing revealed significantly higher scores in the aCSF+6-OHDA (n=6) (**p<0.01) treatment group and ScP+6-OHDA (n=4) (*p<0.05) compared to the TP5+6-OHDA (n=4) treatment group (Figure 2B). Groups not receiving TP5 demonstrated considerable difficulty traversing or complete inability to transverse the narrow beam supporting TP5 protection against balance impairments from 6-OHDA.

**Figure 2.**
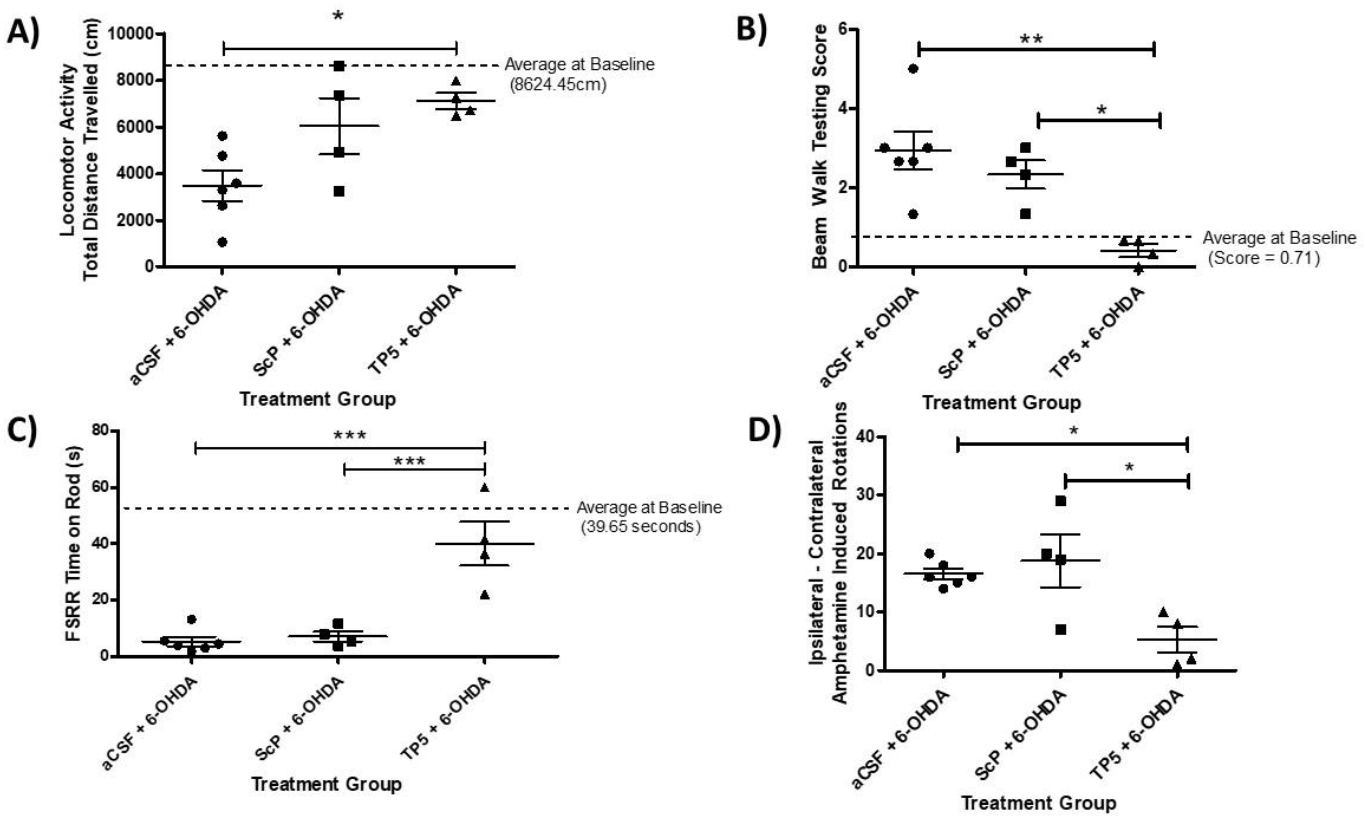
A) Locomotor activity in treated 6-OHDA animals. Baseline data averaged for all animals shown as a dotted line (8624.45cm) for visual representation of “normal” for this cohort. A significant effect of group was found. One Way ANOVA and Tukey’s post hoc testing found aCSF+6-OHDA and TP5+6-OHDA were significantly different **2B) Beam walk scores are lower in TP5 pretreated rats**. Higher scores represent increased difficulty in completing the beam test. Collective baseline average is shown on the graph by a dotted line for reference of a “normal” score (0.71). One Way ANOVA with Tukey’s post hoc test show aCSF and ScP prior to 6-OHDA have increased scores therefore increased difficulty performing the beam test compared to TP5 pretreated animals. **2C) Pretreatment with TP5 has a significant effect on fixed speed rotarod endurance**. The dotted line represents average endurance times of all animals at baseline (39.65 seconds). TP5+6-OHDA animals remained on the rod at 10 rotations per minute, significantly longer than aCSF+6-OHDA or ScP+6-OHDA. **2D) TP5 before 6-OHDA reduced ipsilateral amphetamine induced rotations**. TP5 pretreated animals made significantly less ipsilateral rotations than aCSF and ScP pre-treated animals. Statistical significance is depicted as *p<0.05, **p<0.01, ***p<0.001.

### 3.3 TP5 pre-treated animals do not display significantly reduced endurance using fixed speed rotarod

Fixed speed rotarod (FSRR) highlights gait and coordination impairments. At 10 rotations per minute (rpm), FSRR confirmed significant differences in gait between groups (F(13)=21.39, p=0.0002) using One Way ANOVA. Tukey’s post hoc test revealed a significant difference between aCSF+6-OHDA (n=6) (***p<0.001) animals and ScP+6-OHDA (n=4) (***p<0.001) compared to TP5+6-OHDA (n=4) animals (Figure 2C). TP5 prior to 6-OHDA insult was able to protect rats from developing gait and coordination impairments, as seen through their ability to remain on the rotating rod.

### 3.4 TP5 inhibits amphetamine induced rotations

To phenotypically assess if TP5 protects dopaminergic neurons from 6-OHDA, amphetamine induced rotations were evaluated by considering ipsilateral rotation minus contralateral rotations. One Way - ANOVA found a significant group effect (F(13)=7.298, p=0.0096) and Tukey’s indicated significantly more rotations completed in the aCSF+6-OHDA (n=6) (*p<0.05) and ScP+6-OHDA (n=4) (*p<0.05) group compared to the TP5+6-OHDA (n=4) group over a ten-minute period (Figure 2D). The data phenotypically reveals TP5’s ability to prevent the loss of dopamine neurons acting as a protective agent. This data was reaffirmed using biochemical analysis.

### 3.5 Neuroprotection identified though tyrosine hydroxylase in TP5+6-OHDA treated animals

Motor testing revealed significant impairments in the aCSF+6-OHDA and ScP+6-OHDA groups, while TP5+6-OHDA animals maintained motor function performing close to average baseline values. Thus, dopaminergic protection was investigated indirectly through expression levels of tyrosine hydroxylase (TH) using immunohistochemistry and integrated density values. TH was used as a marker of dopaminergic neurons as this enzyme is a rate limiting step in dopamine synthesis. Immunohistochemical results validated motor impairments found using n=3 per group. One Way ANOVA confirmed an effect of treatment (F(13)=6.073, p=0.0167) and Tukey’s post hoc test showed significantly less TH in the aCSF+6-OHDA and ScP+6-OHDA groups compared to both the Control and TP5+6-OHDA groups (Figure 3). No significant difference was found between control and TP5+6-OHDA treatment groups. The increased TH found in TP5 treated animals suggest potential dopaminergic protection by TP5 however results can only conclude that TH levels are protected.

**Figure 3.**
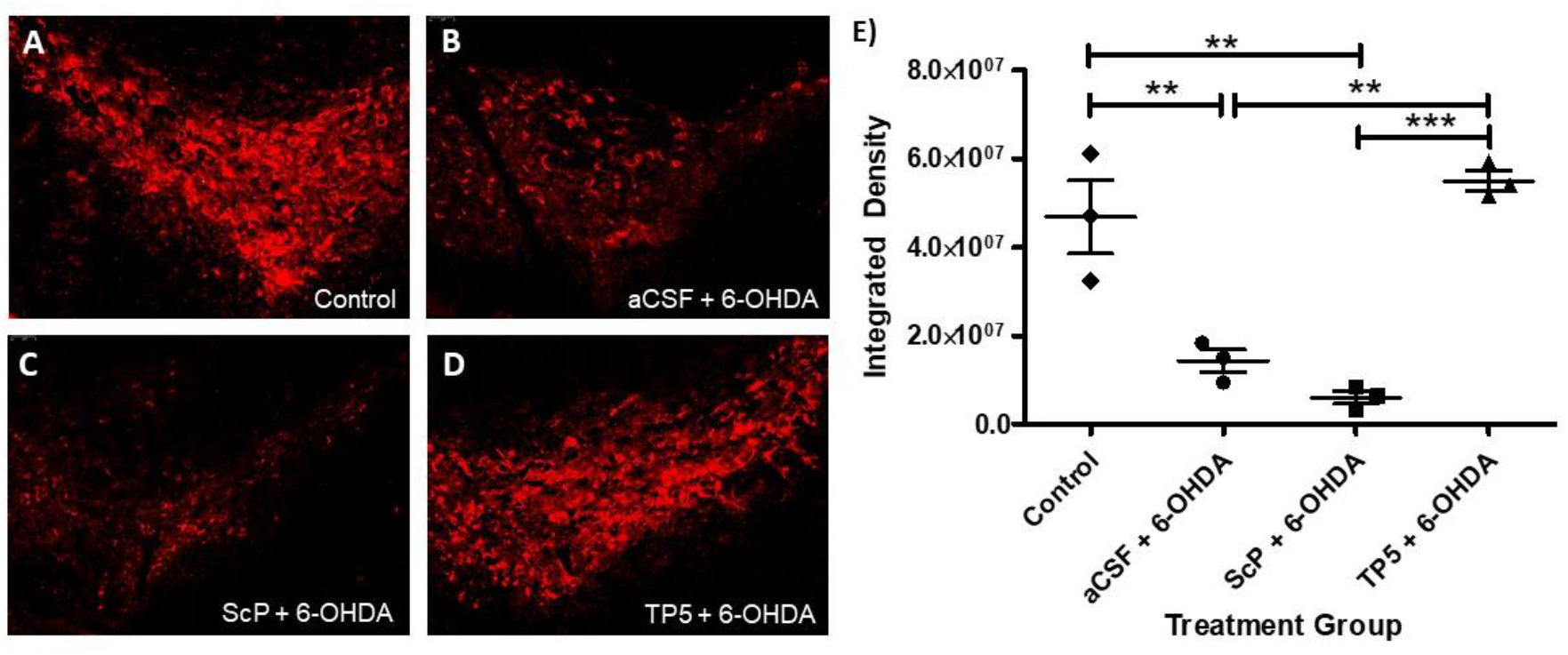
Pre-treatment with TP5 provides protection for tyrosine hydroxylase (TH). TH in the SN was detected using immunohistochemistry. Images shown correspond to the following treatment groups **A)** Healthy control with no treatment **B)** aCSF+6-OHDA **C)** ScP+6-OHDA **D)** TP5+6-OHDA. **E)** Integrated density quantified the presence of TH. Significantly higher integrated density was found in Control and TP5+6-OHDA compared to aCSF+6-OHDA and ScP+6-OHDA (*p<0.05, **p<0.01, ***0.001).

### 3.6 TP5 inhibits hyperactivated CDK5 activity in 6-OHDA lesioned rats

After establishing neuroprotective effects of TP5, the mechanism for neuroprotection was considered. TP5 is known to specifically inhibit CDK5/p25 to reduce CDK5 hyperactivity in pathological states. CDK5 activation in the SN (site of injection) was investigated. All values are expressed as normalized to control, the control being a sham surgery group receiving only aCSF in place of treatment and 6-OHDA. This controlled for volumes administered and potential changes in CDK5 activity produced by surgical intervention. Figure 4 demonstrates a significant effect of treatment using a One Way ANOVA (F(17)=12.11, p=0.0004). Both aCSF+6-OHDA (n=6) (**p<0.001) and ScP+6-OHDA (n=4) (***p<0.001) groups had significantly increased CDK5 activity compared to aCSF only treated controls (n=4). This specifically shows 6-OHDA does hyperactivate CDK5/p25. TP5+6-OHDA did not significantly differ from aCSF treated animals (p>0.05) and was significantly different from ScP+6-OHDA (**p<0.01)TP5 treatment prior to 6-OHDA (n=4) reduced CDK5 hyperactivation at least partially and suggests a mechanism by which TP5 was able to provide neuroprotection from 6-OHDA induced dopaminergic degeneration. One outlier was detected in the aCSF+6-OHDA group using Grubbs outlier test. With removal of this outlier, the result still stands with significance level between ScP+6-OHDA and TP5+6-OHDA after post hoc testing showing more of a significant difference (**p<0.01).

**Figure 4.**
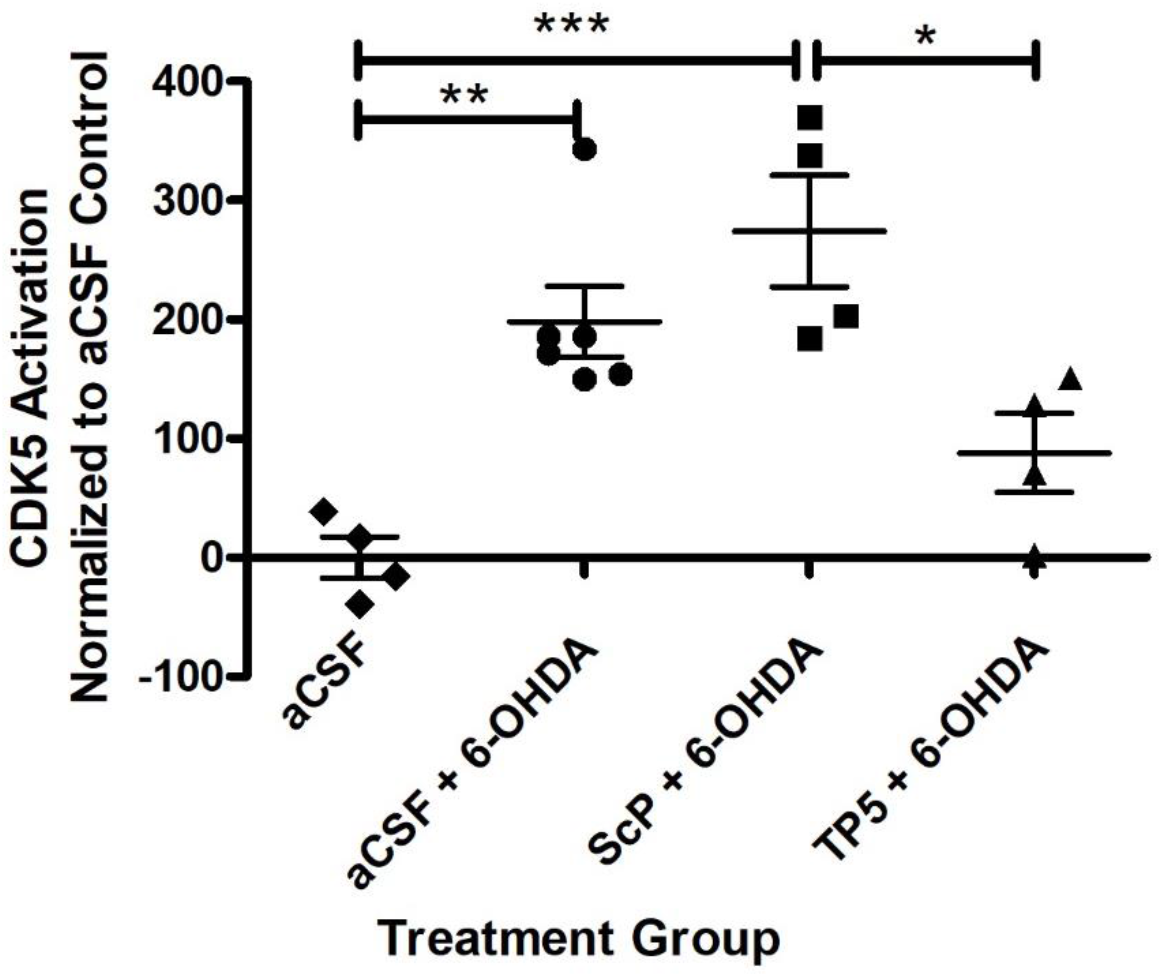
6-OHDA increases CDK5 activity which is reduced by TP5. The graph depicts CDK5 activation based on normalization to aCSF sham surgery controls. Increased CDK5 activation was found in aCSF+6-OHDA and ScP+6-OHDA groups compared to aCSF controls. CDK5 activation was reduced in TP5+6-OHDA compared to ScP-6-OHDA. The TP5+6-OHDA group was not significantly different from aCSF sham surgery animals (*p>0.05, **p<0.01, p<0.0001).

## 4.0 Discussion

The underlying mechanisms for DAergic neuronal loss of PD is not fully understood. Current intervention supplements dopamine for symptomatic treatment but remaining neurons degenerate eliciting the need for neuroprotective agents. The 6-OHDA model produces significant dopaminergic loss and manifests motor abnormalities similar to PD patients, thus was used in the current study to further provide proof of concept evidence and therapeutic potential of TP5 (Binukumar et al., 2014, 2015). Through motor and biochemical evaluation, this study successfully demonstrated treatment with TP5 prior to 6-OHDA significantly acted as a protective intervention. Biochemical analysis discovered CDK5 hyperactivity is increased in the 6-OHDA model and TP5 is able to at least partially reduce this hyperactivation. Immunostaining of higher TH levels in the SN also supports proof of concept dopaminergic neuron protection. Overall, this study demonstrates inhibition of CDK5/p25 is a potential therapeutic target for PD and shows behavioural and biochemical neuroprotective properties of TP5. These findings are consistent with previously reported observations in the MPTP model of PD (Binukumar et al., 2014, 2015)

Oxidative stress is a current hypothesis for dopaminergic degeneration. The neurotoxin 6-OHDA induces oxidative stress (Smith and Cass, 2007). Oxidative stress increases the influx of calcium producing the hyperactivation of CDK5/p25, further increasing oxidative stress (Sun et al., 2008). Calcium dependent enzyme calpain then cleaves p35 (pro-survival promoter) to p25 (pro-apoptotic promoter) favouring the CDK5/p25 complex. This CDK5/p25 complex facilitates apoptotic processes through phosphorylation of p53 (Zhang et al., 2002). In this study we confirmed CDK5/p25 hyperactivation in the 6-OHDA model compared to sham surgery controls, supporting this oxidative stress hypothesis. Interestingly, CDK5 activity was not found to be significantly reduced between aCSF+6-OHDA and TP5+6-OHDA however, we did see significance between ScP+6-OHDA and TP5+6-OHDA. This is interesting because it may indicate the hyperactivation may not be fully overcome by TP5 however reconciling this with behavioural data, this amount of inhibition is still able to produce behavioural benefits. Increased sample size would be able to better make this distinction. Further, TH+ neurons were significantly reduced in ScP+6-OHDA and aCSF+6-OHDA treated animals compared to TP5+6-OHDA animals. Together this suggests CDK5/p25 inhibition through TP5 may provide dopaminergic neuronal protection in the SN.

TP5 is an innovative peptide that takes advantage of promoter specific regulation of CDK5 by selectively inhibiting the CDK5/p25 complex. Current therapeutics aiming to reduce CDK5 activity also inhibit the beneficial effects of CDK5/p35 and lack therapeutic efficacy (Shah and Lahiri, 2014; Siklos et al., 2015). This study supports CDK5/p25 hyperactivity as a viable treatment target and provides evidence for TP5 as a potential therapeutic through behavioural and biochemical results. Behaviourally, motor function and coordination tests provide phenotypic results. The motor tests chosen in this study reflect rigidity, bradykinesia, akinesia, gait abnormalities, and postural imbalances (Taylor et al., 2010). Each of which relate to different PD related motor impairments allowing for translatability. Protective attributes of TP5 were seen in both motor activity and motor function tests as TP5+6-OHDA treated animals performed close to baseline levels using the beam walk, fixed speed rotarod and locomotor chambers. Amphetamine induced rotations was performed to provide a behavioural proxy for dopaminergic protection. Conceptually, unilateral 6-OHDA administration causes dopaminergic neuron loss and receptor super sensitivity on one side. Administration of Amphetamine, a dopamine agonist, will therefore exert effects differently to each hemisphere of the brain and this manifests as rotational behaviour in the rat. Here we demonstrated animals treated with TP5 prior to 6-OHDA did not exhibit this stereotypical behaviour reminiscent of receptor super sensitivity but the 6-OHDA animals without protection did rotate as expected. Together behavioural results support TP5 administered before 6-OHDA protected the against the development of several motor impairments.

Proof of concept studies evaluating hallmark dopaminergic degeneration are imperative for identifying neuroprotective qualities of potential PD therapeutics. This study exemplified protection by TP5 within the SN, supporting further investigation of this peptide and conformationally constrained analoguesthat easily penetrate blood brain barrier. This study however is not without its limitations. More power could be given to the study had larger sample sizes been used and with this proof of concept exhibited follow up work will increase sample sizes. This study also looked at TH as an indirect indicator of DA and thus can only conclude that TH levels are protected, not specifically dopaminergic neurons. While changes in dopamine can be assumed to correlate with changes in TH because TH is the rate limiting step in DA synthesis, we cannot differentiate if this is directly due to the number of DA neurons or functioning of DA neurons. Finally, this study only investigated changes at the SN level and did not consider downstream striatal changes that are involved in motor function. Motor function was impacted as exemplified by several behavioural results however striatal dopamine should be directly considered in later studies. This study provides the proof-of-concept building block for future studies. To continue exploring CDK5/p25 hyperactivation and the potential for TP5, we can investigate TP5 alone in healthy animals. Post mortem studies find CDK5 is co-localized with Lewy bodies suggesting CDK5/p25 plays a role in Lewy body formation (Brion and Couck, 1995; Nakamura et al., 1997; Takahashi et al., 2000) which can also be explored in transgenic model of PD. Finally, non-motor features of PD need to also be considered in drug discovery and design. Non-motor symptoms often go untreated due to the involvement of neurotransmitter systems other than dopamine (Schapira, 2009; Schapira et al., 2017). Therefore, TP5’s affect on non-dopaminergic systems should also be explored for efficacy as well as potential adverse effects.

The use of neurotoxins such as 6-OHDA includes some risk by producing a large amount of toxicity in the aCSF+6-OHDA and ScP+6-OHDA groups. Two of the animals in the study were euthanized due to endpoint from these groups. No animals from the TP5+6-OHDA group reached endpoint.

Given the progressive nature of PD, the need for protective treatments is high and remains to be a significant therapeutic challenge. This study demonstrated CDK5/p25 specific inhibition by TP5 produces neuroprotective effects when used in the 6-OHDA lesion model of PD at phenotypic and molecular levels. This study also further supports the involvement of CDK5/p25 hyperactivity as a possible mechanism in the pathophysiology of PD as its inhibition can offer dopaminergic neuroprotection and protect against motor symptom development. Overall, this study supports further investigation of CDK5/p25 hyperactivity’s involvement in PD pathology and investigation of TP5 as a potential protective therapeutic for neurodegenerative disorders.

## 6.0 Acknowledgments

The authors would like to acknowledge Aurore Latranga for assistance with behavioural tests. Authors also appreciate the support from all Mishra laboratory members in the completion of this work.

